# Isolation and characterization of barley (*Hordeum vulgare*) extracellular vesicles and their role in RNAi-based crop protection

**DOI:** 10.1101/2021.04.26.441409

**Authors:** T Schlemmer, L Weipert, C Preußer, M Hardt, A Möbus, P Barth, T Busche, A Koch

## Abstract

The demonstration that spray-induced gene silencing (SIGS) can confer strong disease resistance bypassing the laborious and time-consuming transgenic expression of double-stranded (ds)RNA to induce gene silencing of pathogenic targets was groundbreaking. However, future field applications will require fundamental mechanistic knowledge on dsRNA uptake, processing, and its transfer. There is increasing evidence that extracellular vesicles (EVs) mediate the transfer of transgene-derived small interfering (si)RNAs in host-induced gene silencing (HIGS) applications. Here, we examined the role of EVs regarding the translocation of sprayed dsRNA from barley (*Hordeum vulgare*) to the target fungus *Fusarium graminearum*. We found barley EVs with 156 nm in size containing predominantly 21 and 19 nucleotide (nt) siRNAs starting with a 5’-terminal Adenine. Notably, barley EVs contain less siRNA compared to EVs isolated from transgenic HIGS Arabidopsis plants. Together our results further underpin mechanistic differences between HIGS and SIGS applications and a minor role of EVs in SIGS.

## Introduction

RNAi-based plant protection strategies represent a powerful tool to address the goals of the European “farm to fork” strategy to reduce the usage of pesticide about 50% till 2030. As an alternative to conventional pesticides, RNAi-based plant protection holds enormous potential to prevent further drastic loss of biodiversity. Over the last two decades, more than 170 studies have demonstrated the feasibility of controlling agronomically and horticulturally relevant plant diseases by utilizing transgenic expression (host-induced gene silencing [HIGS] ^1^) and exogenous application (spray-induced gene silencing [SIGS] ^2^) of double-stranded RNA (dsRNA) to trigger post-transcriptional gene silencing of target genes in various plant pathogens and pests^3^. In addition to the academic proof-of-concept for numerous pathosystems, RNAi technology has further advanced to enable lab-to-field transitions (e.g., costs, risk-assessment, formulations, and regulations). Despite such significant achievements, however, we still lack a mechanistic understanding of these technologies. For example, the mechanisms underlying the transfer and uptake of transgene- or spray-derived RNAs during plant-fungal interactions remain ill-defined, yet they play a pivotal role in determining efficacy and specificity for RNAi-based plant protection. We predict that closing these gaps in knowledge will facilitate the development of novel integrative concepts, precise risk assessment, and tailor-made RNAi therapy for plant diseases. To this end, we assessed the role of extracellular vesicles (EVs) in the transfer of SIGS-inducing RNAs.

Recent data suggest that, analogous to the role of mammalian exosomes in cell-to-cell communication, fungi rely on a bidirectional sRNA transport system via EVs^4^. It has been shown that EVs purified from *Arabidopsis thaliana* leaf extracts and apoplastic fluids contain transgene-derived small interfering RNAs (siRNAs). Furthermore, RNA sequencing (RNA-seq) analysis reveals that EVs from plants expressing CYP3RNA, a 791 nucleotide (nt) long dsRNA originally designed to target the three *CYP51* genes of the fungal pathogen *Fusarium graminearum* (*Fg*), contain CYP3RNA-derived small interfering RNAs (siRNAs) ^5^. Notably, the EVs’ cargo retained the same CYP3RNA-derived siRNA profile as that of the respective leaf extracts, suggesting no selective uptake of specific artificial sRNAs into EVs. Moreover, mutants of the endosomal sorting complex required for transport-III (ESCRT-III) were impaired in HIGS, and EVs were free of CYP3RNA-derived siRNAs^5^. The latter serves as further indication that endosomal vesicle trafficking supports the transfer of transgene-derived siRNAs between donor host cells and recipient fungal cells. Although the number of EV-contained siRNAs was low, we lack information on the minimum concentration of siRNAs required inside an EV to induce HIGS. Notably, we demonstrated previously that *Fg* can take up long dsRNA that is processed by its own RNAi^6,7^, which may explain why we observed greater silencing efficacy of the fungal target genes^7^. Feeding on dead plant tissue, necrotrophic fungi may take up topically-applied dsRNA or dsRNA that was delivered to the xylem^2^. Consequently, we speculate that the role of EVs is minor in the SIGS-*Fg*-barley system. In the present study, we isolated, for the first time, EVs from dsRNA-sprayed barley leaves and analyzed their RNA cargo to determine similarities and differences between EVs’ RNA cargo isolated by HIGS and SIGS strategies, respectively.

## Results and Discussion

To study whether barley (*Hordeum vulgare*) EVs contain SIGS-derived RNAs, we established a protocol for EV isolation from barley leaves by adjusting the EV isolation protocol we had employed for *Arabidopsis* preparation^5^. Accordingly, unsprayed harvested leaf segments were freshly cut on both ends immediately before being submerged in the vesicle isolation buffer. The duration of vacuum infiltration was increased to four min and repeated three times to fully infiltrate the barley leaves. In comparison, *Arabidopsis* leaves required only 1 min per round to achieve full leaf infiltration. To harvest the apoplastic fluid, the centrifugation speed was adapted from 700 xg to 1000 xg for 20 min at 4 °C. Purified barley EVs exhibited a highly diverse size distribution with a mean size of 156 +/− 12.2 nm, which is slightly larger than the mean size of EVs isolated from *Arabidopsis* (139 +/− 7.7 nm^5^, Figs. 1a and 1b) but still fits well within the standard 50–300 nm range for plant EVs^4^. Transmission electron microscopy (TEM) revealed no obvious differences in electron density for barley EVs compared to *Arabidopsis* EVs^5^ (Fig. 1a). Notably, nanoparticle trafficking analysis (NTA) and TEM displayed a strong heterogenicity of size among barley EVs compared to *Arabidopsis* EVs. In addition, NTA revealed several peaks between 100 and 250 nm, which were confirmed by TEM-based size measurements (Figs. 1a and 1b). However, additional verification is required to confirm differences in EV biogenesis between monocot and dicot plant species. To our knowledge, this is the first report on EVs isolated from barley leaves. Thus, we lack an EV marker for immunodetection, which is necessary to prove the EVs’ origin. For *Arabidopsis* EVs, syntaxin PENETRATION1 (PEN1)^8^ and TETRASPANIN8 (TET8)^9^ are referenced EV markers. Currently, the limited information on EV markers in *Arabidopsis* as the plant model organism further impedes efforts to identify possible barley EV markers. Based on amino acid similarity, we located 10 homologs for PEN1 and seven homologs for TET8 in barley (Figs. 1c and 1d).

**Fig. 1.**
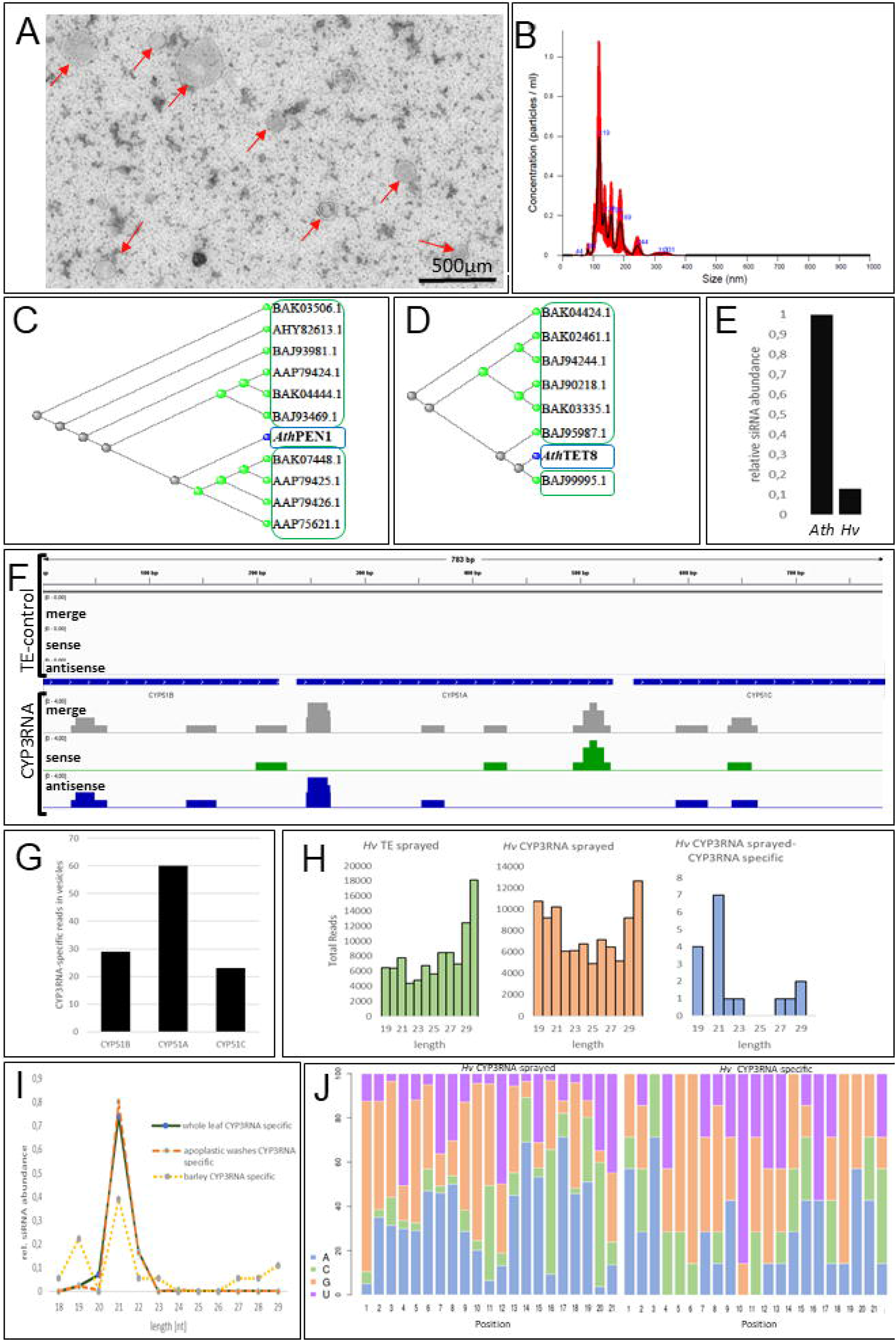
(a) Barley EVs were negatively stained onto cooper formvar meshes using 2% uranyl acetate. (b) Next, 5 µL of purified EVs were diluted up to a volume of 500 µL. Vesicle suspension was loaded into Nanosight NS300 (Malvern Panalytical). Five measurements were performed at 25° C, and size, concentration prediction, and statistical analysis were performed using the NTA 3.2 Dev Build 3.2.16 software. (c)(d) *Arabidopsis thaliana* PEN1 (AT3G11820) and TET8 (AT2G23810) paralogs of *Hordeum vulgare* subsp. vulgare were predicted by the NCBI’s protein BLAST service (https://blast.ncbi.nlm.nih.gov/Blast.cgi). (e) Relative amounts of siRNAs isolated from Arabidopsis and barley EVs. (f) RNA was isolated from mock and dsRNA-treated barley leaves. Indexed sRNA libraries were pooled and sequenced on the Illumina MiSeq Platform (1 x 36 SE). The readings were then mapped onto the CYP3RNA sequence. (g) Number of reads aligned to each CYP3RNA fragment (*CYP51A, CYP51B, CYP51C*) were counted or (h) sorted based on their size. (i) siRNA size distribution of barley EVs were compared with siRNA size distribution of *Arabidopsis* EVs. (j) The nucleotide distribution for every position was counted for the 21 nt long siRNAs with perfect complementarity towards the CYP3RNA precursor.

To verify the role of EV-mediated transport of SIGS-derived siRNA, barley leaves were sprayed with CYP3RNA, as described^2^. EVs were isolated from apoplastic fluids, and EV RNA cargos were examined by RNA-seq. sRNA-profiling of barley EVs revealed fewer CYP3RNA-derived siRNAs (Fig. 1e), as the overall number of siRNAs that mapped to the CYP3RNA precursor was lower than in both *Arabidopsis* samples^5^. Read coverage (number of reads that overlapped at a certain position of the sequence) was also low (Fig. 1f). Notably, the siRNA pattern demonstrated a bias towards siRNAs that matched the middle of the CYP3RNA triple construct (Fig. 1g), which was observed for *Arabidopsis* as well^5^. Further analysis enabled the identification of several of the same siRNAs in both systems, *Arabidopsis*-HIGS and barley-SIGS. Our findings also indicate that the majority of siRNAs are 21 nt in length (Fig. 1h) and preferentially start with an A (Fig. 1j), while siRNAs in EVs isolated from transgene-expressing (HIGS) *Arabidopsis* plants begin with an A or U^5^. Based on sRNA-seq data revealing 5’-identities and lengths of HIGS-derived siRNAs, we can speculate regarding contributing RNA-binding proteins, insofar as they are known for the specific pathosystem. Interestingly, barley EVs revealed a second peak for 19 nt siRNAs, which we did not observe in EVs from *Arabidopsis* (Figs. 1h and 1i). This finding—along with previously discovered differences in efficiencies between dsRNA originating from endogenous expression (HIGS) and dsRNA originating from exogenous spray^10^—underlines mechanistic differences between HIGS and SIGS regarding dsRNA uptake, processing, and transfer. In sum, our current knowledge supports a model of HIGS that involves both plant Dicer-mediated processing of transgene-derived dsRNA into siRNAs and ESCRT-III components mediating RNA transfer—possibly via EVs. Nevertheless, the process by which EVs traverse the plant-fungal interface, as well as the question of whether *Fg* takes up EVs or siRNA/dsRNA that is released from EVs prior to uptake, remains unclear. In contrast, sprayed dsRNA is only partly processed by plant Dicers, while unprocessed dsRNA was shown to be taken up by *Fg*^6,7^. This may explain the lower amounts of siRNA in barley EVs compared to *Arabidopsis* EVs. However, future research must determine whether the loading of unprocessed dsRNA into EVs contributes to SIGS.

When examined holistically, our data suggest that EV loading with CYP3RNA-derived siRNA differs depending on whether HIGS and SIGS strategies are used. The data thus underline mechanistic differences in the uptake and transfer mechanisms of siRNA/(dsRNA). Given these differences, it is necessary to integrate our current knowledge regarding the molecular properties (e.g., pathogen- or pest-specific RNAi mechanisms) with the related strengths and limitations (e.g., routes of dsRNAs and siRNAs) of the chosen pathosystem. This information must be considered when determining which HIGS/SIGS strategy is best.

## Material and Methods

### Plant cultivation and CYP3RNA spray-application

160 second leaves of barley cv. Golden Promise were harvested from plants growing for 3 weeks under long day conditions (16 h light, 22°C, 60% humidity). The leaves were transferred to square petri dishes with 1% agar and divided into two groups. The upper part of the first group were sprayed with CYP3RNA diluted in TE buffer and the second group was sprayed with TE buffer as mock control as previously described^2^ and incubated for 48 h before EV isolation was performed.

### Negative staining and transmission electron microscopy (TEM)

For TEM, copper formvar-coated 300-mesh electron microscopy grids were glow discharged prior to sample application for 40 sec. Subsequently, 5 µl of the sample resuspended in PBS were applied to the grids. Samples were dabbed off using Whatman filter paper and grids were washed three times in 50 µl of 2% uranyl acetate and one time with distilled water. Needless staining or fixing solutions, buffers and water were removed by Whatman paper between each step. Finally, grids were air dried. Preparations were inspected at 120 kV under zero-loss conditions (ZEISS EM912a/b) and images were recorded at slight underfocus using a cooled 2k x 2k slow-scan ccd camera (SharpEye / TRS) and the iTEM software package (Olympus-SIS).

### Vesicle size and concentration measurements by nanoparticle trafficking analysis (NTA)

For size and concentration prediction, purified barley EVs were diluted (1:50) with PBS. Subsequently, 500 µL of vesicle suspension was loaded into Nanosight NS300 (Malvern Panalytical). 5 measurements were performed at 25°C and size, concentration prediction and statistical analysis were performed by the NTA 3.2 Dev Build 3.2.16 software.

### Determine siRNAs originating from CYP3RNA

Vesicle RNA was isolated using the Single Cell RNA Purification Kit (Norgen Biotek) according to the manufacturer’s instructions described for cells growing in suspension. RNA concentrations were determined using the NanoDrop spectrophotometer (Thermo Fisher Scientific) and RNA was stored at -80°C before sending samples to RNA sequencing. Indexed sRNA libraries were constructed from RNA isolated from vesicles with the TruSeq ^®^ Small RNA Library Prep Kit (Illumina) according to the manufacturer’s instructions. Indexed sRNA libraries were pooled and sequenced on the Illumina MiSeq Platform (1 x 36 SE) and the sequences were sorted into individual datasets based on the unique indices of each sRNA library. The quality of the datasets was examined with FastQC (version 0.11.9) (https://www.bioinformatics.babraham.ac.uk/projects/fastqc/) before and after trimming. The adapters were trimmed using cutadapt (version 2.8)^11^. To filter out bacterial contaminations kraken2 (version 2.1.1)^12^ was used with the database obtained from the MGX metagenomics application^13^. All reads marked as unclassified were considered to be of non-bacterial origin and used for the subsequent alignment. The trimmed and filtered reads were mapped to the CYP3RNA sequence using bowtie2 (version 2.3.2) ^14^ to identify siRNAs derived from the precursor dsRNA sequence. The mappings were first converted into bedgraph using bedtools (version 2.26.0)^15^ and then to bigwig using bedGraphToBigWig^16^. These files were used for visualization with IGV^17^. Read coverage is defined as the number of reads that match at a certain position of the sequence.

### Determine frequency of different RNA species

To determine RNA species, reference genome and annotation of Hordeum Vulgare (IBSC v2 – release 47) were downloaded from EnsemblPlants^18^. Adapter trimming of raw reads was done with TrimGalore (version 0.6.4) (https://www.bioinformatics.babraham.ac.uk/projects/trim_galore/) which used cutadapt (version 2.8)^11^. In this process all reads which became shorter than 18 nt were filtered out. Afterwards, nucleotides with a phred score below 20 and reads retaining less than 90% of their nucleotides in this process were removed using FASTQ Quality Filter from the FASTX-toolkit (version 0.0.14) (https://github.com/agordon/fastx_toolkit). The bacterial contaminations were filtered out as demonstrated in the previous section. The remaining reads were aligned to the reference genome using STAR (version 2.7.3a) ^19^. The number of different RNA species was examined in R (version 4.0.2) (R Core Team, 2020) using featureCounts from the package Rsubread (version 2.2.5)^20^. featureCounts was run for each sample using the previously downloaded annotations of Arabidopsis. Following RNA types were examined: “lncRNA”, “pre_miRNA”, “mRNA”, “ncRNA_gene”, “rRNA”, “snoRNA”, “snRNA” and “tRNA”. All alignments that could not be assigned to a feature were considered as “not assigned”.

## Author Contributions

T.S., L.W. and A.K. wrote the manuscript; A.K. and T.S. designed the study; T.S., and L.W., conducted the experiments; T.S., L.W., A.K., and P.B. analyzed all data and drafted the figures. T.B. conducted RNA-seq experiments and T.B., and P.B. performed bioinformatics analysis. All authors reviewed the final manuscript.

## Acknowledgements

We thank C. Birkenstock and U. Schnepp for excellent plant. cultivation and M.Sc. C. Pfafenrot and M.Sc. M. Mosbach for helping with the NTA measurements. This work was supported by the Deutsche Forschungsgemeinschaft, Research Training Group (RTG) 2355 (project number 325443116) to AK and TS. We acknowledge access to compute resources of the Bielefeld-Gießen Center for Microbial Bioinformatics (BiGi) financially supported by the BMBF grant FKZ 031A533 within the de.NBI network.

